# Comparison of Deep Learning Tools for Optic Nerve Axon Quantification Finds Limited Generalizability on Independent Validation

**DOI:** 10.64898/2026.03.11.710915

**Authors:** Benton Chuter, Noah Emmert, Min Young Kim, Nikhil Dave, Jay Herrin, Zirui Zhou, Gideon Wall, Abraham Palmer, Hao Chen, T.J. Hollingsworth, Monica M. Jablonski

**Author notes:** **Corresponding Authors:** Benton Chuter, MD, MS & Monica M. Jablonski, PhD, FARVO, Hamilton Eye Institute, 930 Madison Avenue, Memphis, TN 38163.

## Abstract

**Purpose:** Machine learning approaches for automated quantification of optic nerve histology have emerged as potential tools for objective assessment of axonal injury in experimental glaucoma models. However, the generalizability of these models to independent datasets remains unclear. Guided by a scoping review of the literature, this study performed independent validation testing of publicly available models on a novel rat optic nerve dataset to assess their generalizability.

**Methods:** We conducted a scoping review following PRISMA-ScR guidelines. PubMed, EMBASE, Scopus, and Cochrane CENTRAL were searched from 2000 through 2025. Two reviewers independently screened records and extracted data on model characteristics and performance metrics. Additionally, we performed independent validation of three models (AxoNet, AxonDeepSeg, AxoNet 2.0) on a novel rat optic nerve dataset comprising 57 images with 9,514 manually annotated axons. Because AxonDeep is not publicly available, we instead evaluated AxonDeepSeg, a separate publicly available deep learning-based tool that, while not previously applied to optic nerve tissue, is widely used for nerve fiber segmentation.

**Results:** From 2,036 records, four manuscripts describing three deep learning models met inclusion criteria. Published correlation coefficients between model predictions and reference counts ranged from 0.959 to 0.99. On independent validation, performance was reduced: AxoNet 2.0 achieved the highest correlation (r = 0.89), followed by AxonDeepSeg (r = 0.86) and AxoNet (r = 0.79). Segmentation quality metrics revealed high precision (>0.94) but low recall (0.18 to 0.27), with Dice coefficients of 0.29 to 0.40, substantially below published benchmarks of 0.81.

**Conclusions:** Deep learning models for optic nerve histology demonstrate strong within-study performance but show meaningful performance decrements when applied to independent datasets. The observed generalizability gap (correlations 0.07 to 0.182 points below published values) demonstrates the need for standardized validation datasets and multi-center testing before widespread adoption of these tools.

## INTRODUCTION

Retinal ganglion cell (RGC) loss represents the defining pathological feature of glaucoma and other optic neuropathies, conditions that together constitute leading causes of irreversible blindness worldwide (Almasieh et al., 2012). Histological quantification of optic nerve axons provides a direct measure of RGC survival and remains essential for evaluating neuroprotective interventions in experimental models. However, manual axon counting is labor-intensive, subject to inter-observer variability, and impractical for large sample sizes often required in preclinical studies (Nuschke et al., 2015; Smith et al., 2017).

To address these limitations, several approaches have been developed to improve efficiency and reproducibility of axon quantification. Sampling-based approaches that extrapolate from counted subregions introduce additional variability, while fully manual counts of entire nerve cross-sections require hours per specimen (Teixeira et al., 2014; Reynaud et al., 2012). Semi-automated tools such as AxonJ have improved throughput but still require significant operator input and parameter tuning (Zarei et al., 2016).

Deep learning has transformed digital pathology across medical disciplines, with convolutional neural networks (CNNs) achieving expert-level performance in tumor detection, tissue segmentation, and prognostic classification (Echle et al., 2021; Litjens et al., 2016; Srinidhi et al., 2021). These methods learn hierarchical feature representations directly from image data, potentially capturing subtle patterns that manual analysis may fail to detect. In ophthalmology, deep learning applications span retinal disease screening, glaucoma detection from fundus photographs, and optical coherence tomography interpretation (Wang et al., 2019).

Despite this progress, machine learning approaches for optic nerve histology have been described in only a limited number of studies, each employing different model architectures, species, staining methods, and outcome measures. This heterogeneity complicates comparison of model performance and hinders assessment of which approaches may be most suitable for specific applications. Importantly, the generalizability of these models when applied outside their original training environments remains largely untested. Domain shift, wherein models encounter data distributions differing from their training sets, represents a well-documented challenge in medical imaging that can substantially degrade performance (Guan & Liu, 2022; Zech et al., 2018).

This study evaluated the generalizability of published machine learning models for optic nerve axon quantification through independent validation testing guided by a scoping review of the literature. Our objectives were to: (1) identify published models and summarize their architectures and training data; (2) synthesize reported performance metrics across studies; (3) identify gaps in the current evidence base; and (4) evaluate model performance on an independent dataset not used in prior training or validation.

## METHODS

### Scoping Review Design

The scoping review was conducted in accordance with the Preferred Reporting Items for Systematic Reviews and Meta-Analyses extension for Scoping Reviews (PRISMA-ScR) guidelines (Tricco et al., 2018). A scoping review methodology was selected given the anticipated heterogeneity across studies and our objective of mapping the breadth of available evidence rather than synthesizing pooled effect estimates. The protocol was registered with the International Prospective Register of Systemic Reviews (PROSPERO) prior to screening (Registration: CRD420251250152).

### Eligibility Criteria

Studies were eligible for inclusion if they met the following criteria: (1) involved human or animal optic nerve tissue, retinal ganglion cells, axons, myelin, or associated glial cells; (2) applied machine learning methods for quantification, segmentation, or morphometric analysis of histological images; (3) reported quantitative performance metrics comparing automated outputs to reference standards; and (4) were published as peer-reviewed original research articles written in English between 2000 and 2025 (**Supplementary Table S1**).

Studies were excluded if they: did not involve optic nerve or related structures; used only traditional histologic grading without machine learning components; lacked quantitative outcomes; or were reviews, editorials, conference abstracts without full data, or non-peer-reviewed publications.

### Information Sources and Search Strategy

Four electronic databases were searched: PubMed; EMBASE; Scopus; and Cochrane CENTRAL. The search strategy combined three concept blocks using Boolean operators. The first block captured anatomical terms (optic nerve, retinal ganglion cell, axon, myelin, astrocyte, microglia, oligodendrocyte, glia, fibroblast). The second block captured computational methods (deep learning, machine learning, convolutional neural network, CNN, segmentation, automated). The third block captured histological analysis (histology, histopathology, histomorphometry, morphometry, quantification, grading). The complete search strategies for each database are provided in **Supplementary Table S2**.

### Selection Process

Two reviewers independently screened titles and abstracts against predefined eligibility criteria. Records deemed potentially relevant by either reviewer advanced to full-text review. Full-text articles were independently assessed by both reviewers, with disagreements resolved through discussion or adjudication by a third reviewer. The selection process was documented using the PRISMA flow diagram (**Figure 1**).

**Figure 1.**
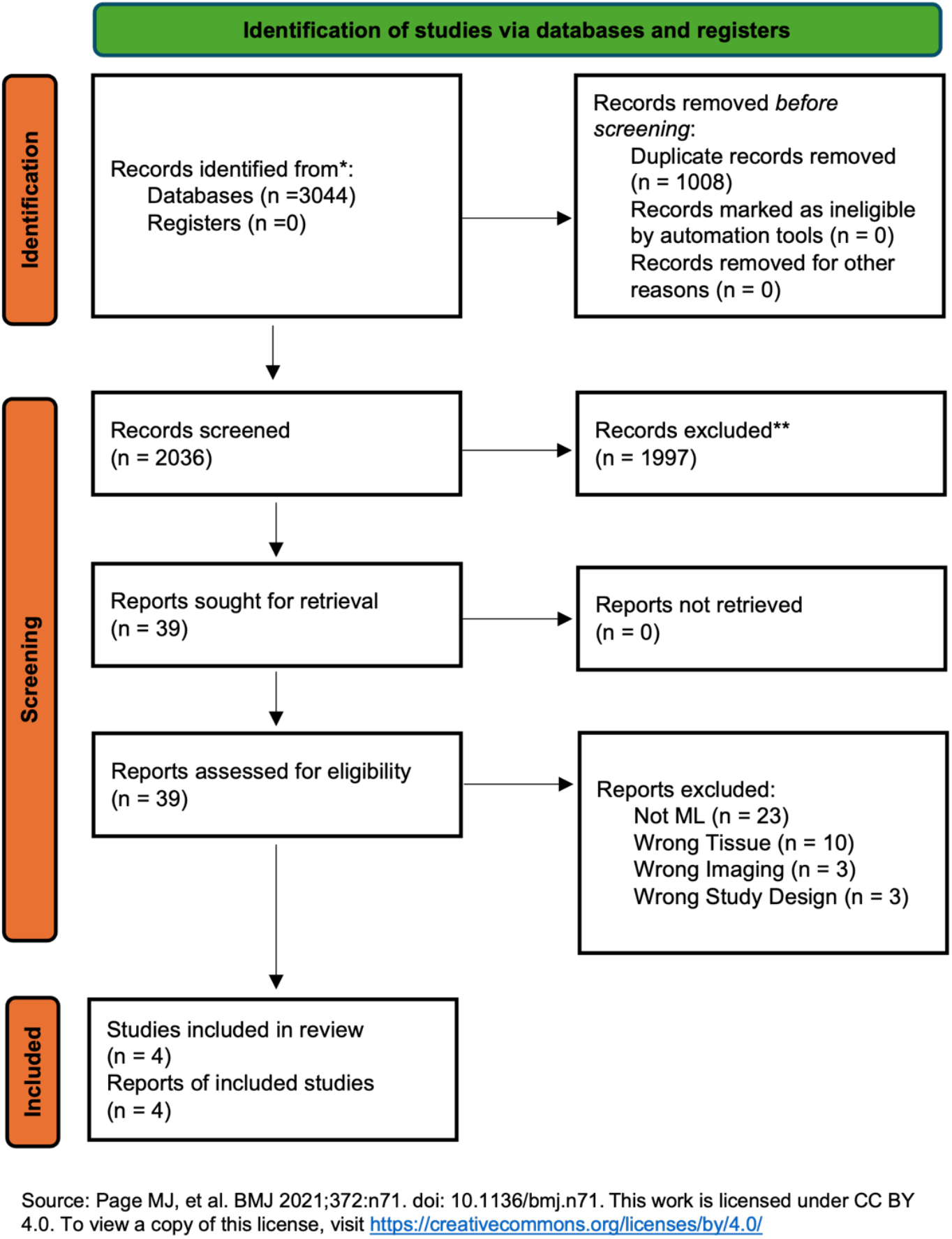
PRISMA 2020 Flow Diagram. Flow diagram outlines the identification, screening, and selection process for the scoping review. Database searches of PubMed, EMBASE, Scopus, and Cochrane CENTRAL identified 2,036 records. After removal of duplicates and title/abstract screening, 1,997 records were excluded. Full-text review of the remaining 39 articles led to exclusion of 35 studies: 18 did not apply machine learning methods; 9 did not involve optic nerve tissue; 5 lacked quantitative performance metrics; and 3 were review articles. Four manuscripts describing three distinct deep learning models (AxoNet, AxonDeep, and AxoNet 2.0) met all inclusion criteria and were included in the final synthesis.

### Data Extraction

A standardized data extraction form was developed and piloted on two studies prior to full extraction. Two reviewers independently extracted data from each included study, with discrepancies resolved by consensus. Complete extracted data are provided in **Supplementary Table S3**. Extracted variables included: study characteristics (authors, year, institution); model characteristics (name, architecture type, training approach); dataset characteristics (species, strain, disease model, staining method, sample sizes); and performance metrics (correlation coefficients, Dice coefficients, mean absolute error, root mean squared error, Bland-Altman limits of agreement).

### Independent Validation Study

To assess model generalizability beyond published performance metrics, we conducted independent validation testing of available models on a novel dataset. The validation dataset comprised 57 paraphenylenediamine (PPD)-stained rat optic nerve cross-section images (256 × 256 pixels, 0.055 micrometers per pixel) containing 9,514 manually annotated axons that were prepared using our previously described methods (Stiemke et al., 2020). Axon annotations were performed by a trained observer who identified and segmented individual axon profiles, distinguishing live from necrotic axons (both categories were combined for evaluation). Annotated axon counts were reported as mean ± SEM, which was used as our ground truth dataset.

Three models were evaluated: AxoNet (using the final_resampled_3-22-2020.hdf5 checkpoint), AxonDeepSeg (using the model_seg_generalist_BF_light configuration; Zaimi et al., 2018), and AxoNet 2.0 (using the standard U-Net architecture). Because AxonDeep (Deng et al., 2021) is not publicly available, we tested AxonDeepSeg as a publicly available alternative for neural network-based axon segmentation. AxonDeepSeg was developed for general axon and myelin segmentation from microscopy data and represents a distinct implementation from AxonDeep, though both employ deep learning for axon identification. All models were applied using their published implementations and default parameters without any fine-tuning or adaptation to the validation dataset.

Performance was assessed at two levels. For axon count agreement, we computed Pearson correlation coefficients, mean absolute error (MAE), and root mean squared error (RMSE) between model-predicted and ground truth axon counts for each image. For segmentation quality, we computed pixel-level metrics including Dice coefficient, intersection over union (IoU), precision, and recall by comparing predicted segmentation masks to ground truth annotations.

### Data Synthesis

Given the scoping nature of the review and the heterogeneity across studies, a narrative synthesis approach was employed for published results. Study and model characteristics were summarized descriptively in tables, and performance metrics were presented graphically. For metrics reported by multiple studies, weighted means were calculated using sample size weights. Independent validation results were analyzed separately and compared to published benchmarks to quantify generalizability gaps.

## RESULTS

### Study Selection

The database search identified 2,036 records across PubMed, EMBASE, Scopus, and Cochrane CENTRAL. After removal of duplicates, 2,036 unique records underwent title and abstract screening. Of these, 1,997 were excluded as clearly not meeting eligibility criteria. The remaining 39 articles underwent full-text review, of which 35 were excluded: 18 did not apply machine learning methods; 9 did not involve optic nerve tissue; 5 lacked quantitative performance metrics; and 3 were review articles. Four manuscripts describing three distinct deep learning models met all inclusion criteria and were included in the final synthesis (**Figure 1**).

### Characteristics of Included Studies

The included studies were published between 2020 and 2023, originating from research groups in the United States. All studies focused on automated quantification of retinal ganglion cell axons in optic nerve cross-sections from experimental glaucoma models. **Table 1** summarizes model characteristics and presents published performance metrics.

**Table 1.** Characteristics and Reported Performance Metrics of Deep Learning Models for Optic Nerve Axon Quantification.

| Model | Author/<br>Year | Architecture | Training | Staining | Species | Sample Size | Reported<br>Correlation | Dice | MAE | RMSE |
| --- | --- | --- | --- | --- | --- | --- | --- | --- | --- | --- |
| AxoNet | Ritch et al.,<br>2020 | U-Net<br>(density) | Supervised | Toluidine<br>blue | Brown<br>Norway rat | 1,514<br>sub-images | $R^2 = 0.938$ | – | 4.4 | – |
| | | | | PPD | NHP* | 494<br>sub-images | $R^2 = 0.944$ | – | 17.7 | – |
| AxonDeep | Deng et al.,<br>2021 | GAN-based<br>FCN | Semi-supervised | PPD | Mouse<br>(DBA/2J,<br>D2.*Lyst*,<br>C57BL/6J<br>and J:DO) | 78<br>sub-images<br>(56 optic<br>nerves) | $R = 0.97$ | 0.81 | 4.4%<br>** | – |
| AxoNet 2.0 | Goyal et al.,<br>2023a,<br>2023 b | U-Net<br>(refined) | Supervised | Toluidine<br>blue | Brown<br>Norway rat | 1421<br>sub-images | $R^2 = 0.92$ | 0.81 | – | 6.18 |
| | | | | PPD | DBA/2J<br>Mouse*** | 22<br>sub-images | $R^2 = 0.98$ | – | – | 63.16 |
| | | | | | NHP* | 50<br>sub-images | $R^2 = 0.97$ | – | – | 74.71 |

*Abbreviations:* GAN = generative-adversarial-network framework; FCN = fully convolutional network; PPD = p-paraphenylenediamine; NHP = non-human primate; DO = Diversity Outbred; MAE = mean absolute error (axons per sub-image); RMSE = root mean squared error. Correlation values represent $R^2$ for AxoNet models and Pearson R for AxonDeep.
\*Species not explicitly named in Ritch et al. (2020) or Goyal et al. (2023b); NHP datasets were annotated from Reynaud et al. (2012) from rhesus and cynomolgus macaques from the Burgoyne Lab (Devers Eye Institute).
\*\*Mean absolute percentage error was reported in this study.
\*\*\*Mouse dataset was annotated from Deng et al. (2021).

AxoNet was described by Ritch et al. (2020) and evaluated on two datasets: a rat dataset comprising 27 optic nerves from 14 Brown Norway rats with unilateral experimental glaucoma (1% toluidine blue staining; 1,514 annotated sub-images), and a non-human primate (NHP) dataset from Reynaud et al. (2012) consisting of optic nerves from rhesus and cynomolgus macaques (paraphenylenediamine [PPD] staining; 494 annotated sub-images).

AxonDeep was described by Deng et al. (2021) and applied to optic nerves from 56 mice across three strains: DBA/2J, D2.*Lyst*, C57BL with blast-induced traumatic brain injury (C57BL/6J), and Diversity Outbred mice (C57BL/J:DO). The dataset included 78 sub-images with PPD staining.

AxoNet 2.0 was described in two publications by Goyal et al. (2023a, 2023b), evaluating the model on Brown Norway rats with unilateral ocular hypertension. The primary dataset included 46 eyes from 23 rats (1% toluidine blue staining; 1,421 annotated sub-images). The model was also validated on DBA/2J mice with blast-induced traumatic brain injury and on rhesus and cynomolgus macaque optic nerves, demonstrating cross-species applicability.

### Model Architectures

All three models employed convolutional neural network architectures based on encoder-decoder designs (**Table 1)**. AxoNet uses a U-Net style architecture that outputs pixelwise axon count density estimates rather than binary segmentation masks. AxonDeep employs a semi-supervised learning framework that integrates a fully convolutional segmentation network with a discriminator network in a generative adversarial network (GAN) architecture. In this approach, the discriminator learns to distinguish model-generated segmentations from expert annotations, providing an additional training signal that reduces the need for fully labeled data. AxoNet 2.0 refined the original AxoNet architecture with improved training procedures and data augmentation strategies. Because the original AxonDeep model was not publicly available, independent validation was performed using AxonDeepSeg (Zaimi et al., 2018), a related U-Net-based architecture for axon and myelin segmentation.

### Published Performance Metrics

Reported performance metrics varied across studies, with correlation coefficients being the most consistently reported measure (**Table 1, Figure 2**). All three models demonstrated strong agreement with manual reference counts in their original publications. For our study, published correlation coefficients (r or r squared) across six model-datasets were converted to r for comparison, which ranged from 0.959 to 0.99.

**Figure 2.**
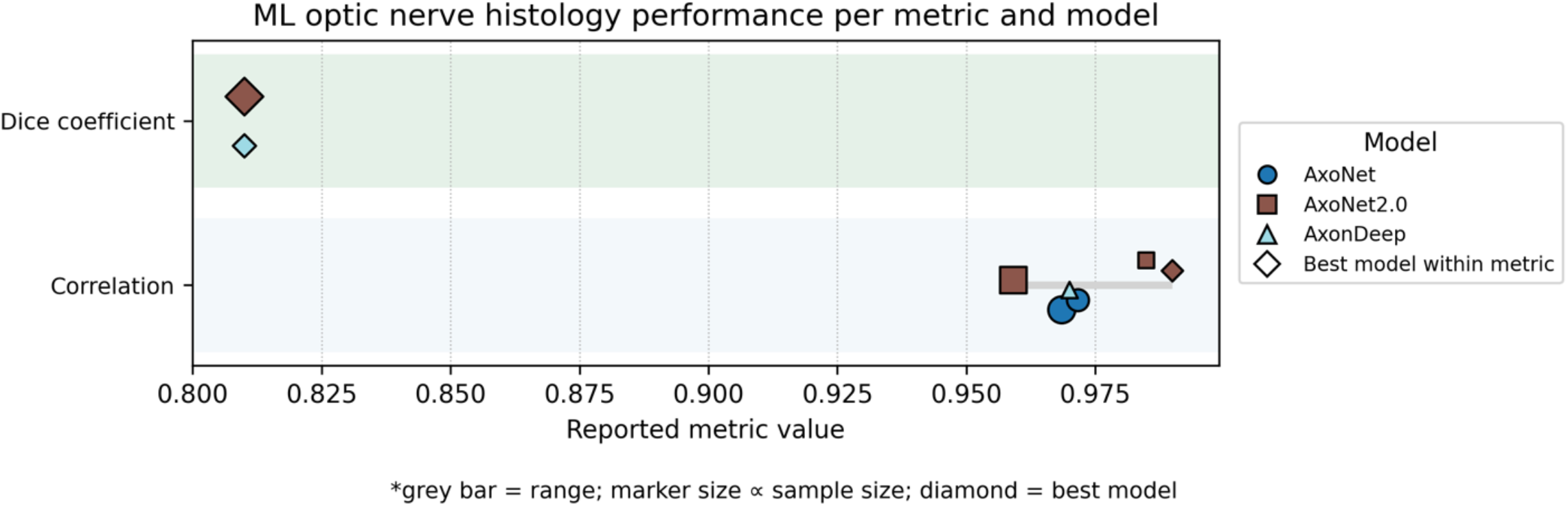
Published Performance Metrics Forest Plot. The forest plot displays reported performance metrics across included deep learning models for optic nerve axon histology. Horizontal axis presents reported Pearson’s r values. Grey bar indicates the range of values across model-dataset combinations; where no grey bar is visible, the range of published values across model-dataset combinations was zero (i.e., identical values were reported). Marker size is proportional to sample size. Diamond markers indicate the best-performing model within each metric category. Correlation coefficients clustered between 0.96 and 0.99, with Dice coefficients of 0.81 reported by models providing segmentation outputs.

AxoNet achieved r = 0.97 on the rat testing subset and r = 0.97 on the NHP testing subset. AxonDeep reported r = 0.97 between semi-supervised predictions and reference counts. AxoNet 2.0 achieved the highest published correlation with r = 0.99 on mouse optic nerves.

Segmentation accuracy was reported as Dice coefficient by two models. Both AxonDeep and AxoNet 2.0 reported Dice coefficients of 0.81, indicating substantial overlap between predicted and reference segmentation masks.

Mean absolute error was reported only for AxoNet: 4.4 axons per sub-image on the rat testing subset and 17.7 axons per sub-image on the NHP testing subset. AxonDeep reported mean absolute percentage error of 4.4% with mouse. The mean squared error was reported for AxoNet 2.0 with 6.18, 63.16, and 74.71 axons for rat, mouse, and NHP models respectively.

### Independent Validation Results

Independent validation across deep learning models on the novel rat optic nerve dataset with known manual axon counts (n = 57 images, 9,514 axons) revealed substantial performance differences from published benchmarks (**Table 2, Figure 3**). Because AxonDeep (Deng et al., 2021) is not publicly available, we evaluated AxonDeepSeg (Zaimi et al., 2018) as a publicly available alternative; direct comparison of AxonDeepSeg results to AxonDeep published metrics should therefore be interpreted with caution.

**Table 2.** Independent Validation Results.

| **Model** | **Correlation (r)** | **MAE** | **RMSE** | **Dice (Ind.)** | **Dice (Pub.)** | **Gap ($\Delta$)** | **IoU** | **Precision** | **Recall** |
| --- | --- | --- | --- | --- | --- | --- | --- | --- | --- |
| AxoNet 2.0 | 0.89 | 69.1 | 81.5 | 0.40 | 0.81 | 0.41 | 0.26 | 0.94 | 0.27 |
| AxonDeepSeg | 0.86 | 77.3 | 93.0 | 0.29 | 0.81 | 0.52 | 0.18 | 0.95 | 0.18 |
| AxoNet | 0.79 | 119.3 | 138.1 | - | N/A | N/A | - | - | - |

Gap represents the difference between published and independent validation performance.
AxoNet outputs density estimates rather than segmentation masks, so segmentation metrics are not applicable. *Abbreviations*: MAE = mean absolute error (axons); RMSE = root mean squared error; Ind. = independent validation; Pub. = published; IoU = intersection over union.

**Figure 3.**
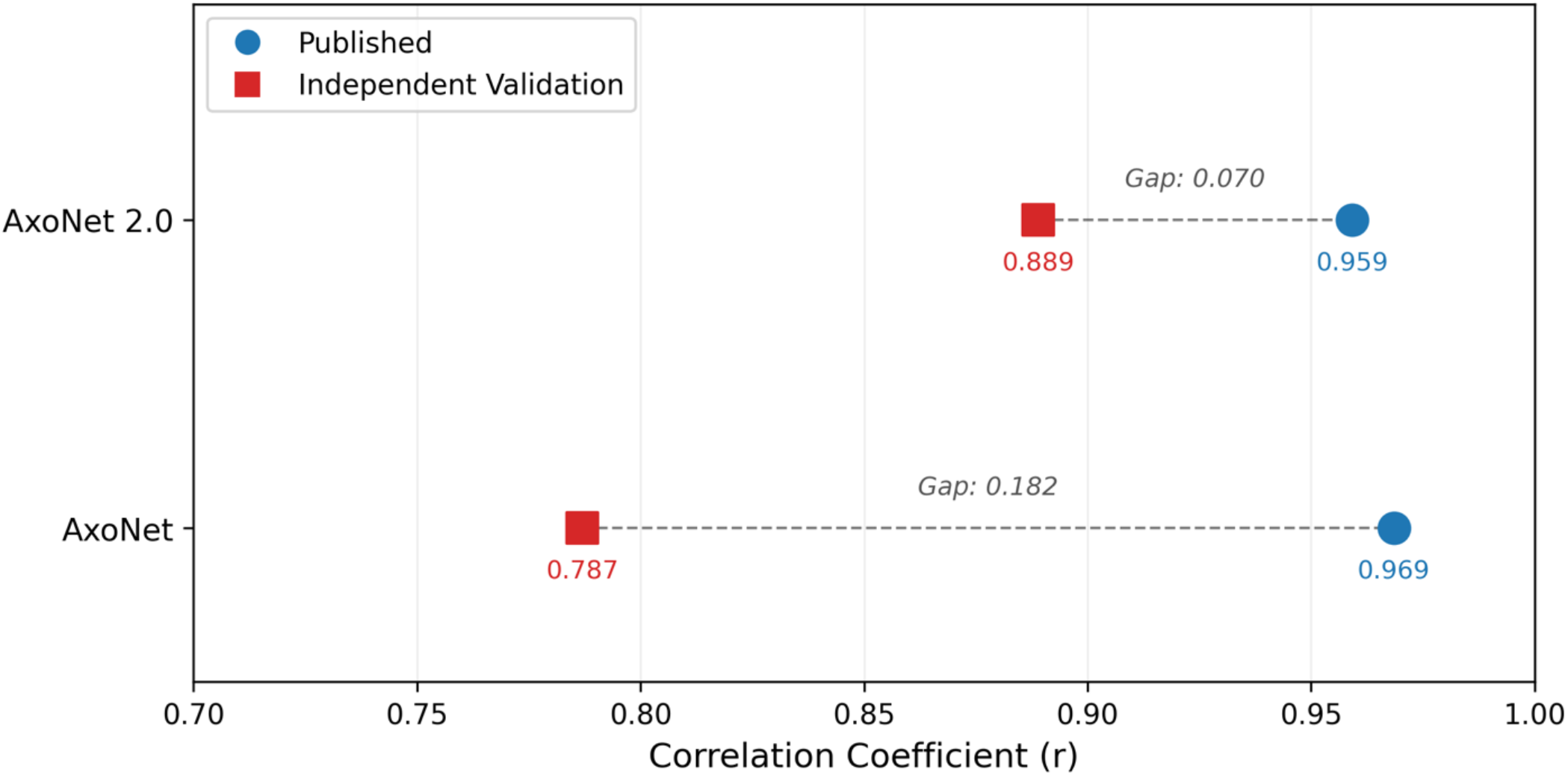
Published vs. Independent Validation Correlation with Reference Axon Counts. Published and independently validated correlation coefficients between model-predicted and reference axon counts are compared. Blue circles indicate values reported in original publications, and red squares indicate performance on an independent dataset of 57 rat optic nerve cross-sections from the authors’ laboratory that were not used in any model’s original training or evaluation. Dashed lines connect paired values, with gap magnitude indicated. AxonDeepSeg is not shown because its original publication reported only segmentation metrics and did not include a correlation benchmark. Performance degradation for models with published benchmarks ranged from 0.070 to 0.182 correlation points.

#### Axon Count Agreement

All models maintained positive correlations with ground truth axon counts, though at reduced levels compared to published values. AxoNet 2.0 achieved the highest correlation (r = 0.89, MAE = 69.1 axons, RMSE = 81.5 axons), followed by AxonDeepSeg (r = 0.86, MAE = 77.3 axons, RMSE = 93.0 axons) and AxoNet (r = 0.787, MAE = 119.3 axons, RMSE = 138.1 axons). The correlation decrements from published values ranged from 0.07 points (AxoNet 2.0) to 0.18 points (AxoNet).

#### Segmentation Quality

Pixel-level segmentation metrics revealed a consistent pattern across models: high precision but low recall **(Table 2)**. AxoNet 2.0 achieved Dice = 0.40, IoU = 0.26, precision = 0.94, and recall = 0.27. AxonDeepSeg achieved Dice = 0.29, IoU = 0.18, precision = 0.95, and recall = 0.18. These Dice coefficients were substantially lower than the published benchmark of 0.81, representing decrements of 0.41 to 0.52 points.

The high precision values (>0.94) indicate that when models identified pixels as belonging to axons, they were usually correct. However, the low recall values (0.18 to 0.27) indicate that models detected only a fraction of the total axon area present in ground truth annotations (**Figure 4)**.

**Figure 4.**
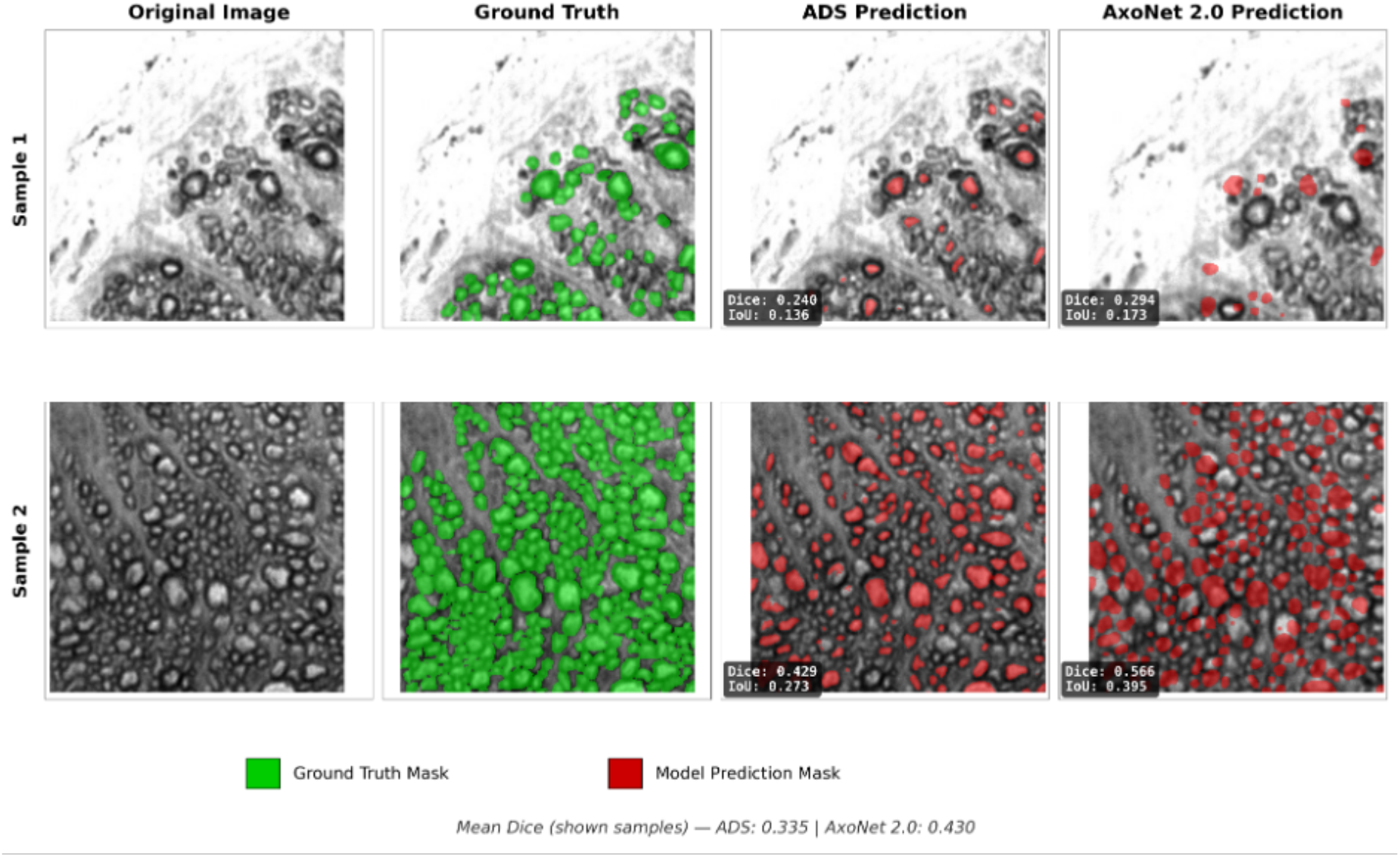
Representative Examples of Axon Segmentation. Representative examples of AxonDeepSeg (ADS) and AxoNet 2.0 axon segmentation on independent rat optic nerve tests are shown.

### Findings

Comparison of published and independent validation results reveals a consistent generalizability gap (**Figure 3**). Published correlation coefficients clustered between 0.96 and 0.97, while independent validation correlations ranged from 0.79 to 0.89. The magnitude of this gap varied by model, with AxoNet 2.0 showing the smallest performance decrement and AxoNet the largest. A detailed comparison of published versus independent validation metrics is provided in **Supplementary Figure S1**.

Notably, model rankings differed between published results and independent validation. AxoNet achieved the highest published correlation for rat models (r = 0.97) but had the lowest correlation in independent validation (r = 0.79) among the three models. Although AxonDeep reported a similar published correlation (r = 0.97), it could not be directly evaluated because its implementation is not publicly available; instead, the alternative model AxonDeepSeg achieved r = 0.86 on independent validation. In contrast, AxoNet 2.0 reported a lower published correlation for rat models (r = 0.96) but achieved the highest correlation in independent validation performance (r = 0.89).

## DISCUSSION

In this study, we identified three deep learning models for optic nerve axon histology quantification through a scoping review and conducted the first independent validation to compare their performance on a novel dataset. Although all models demonstrated strong agreement with reference standards in their original publications —correlation coefficients exceeding 0.96—independent validation revealed meaningful performance decrements ranging from 0.07 to 0.18 correlation points. These findings have important implications for the adoption and further development of automated optic nerve histology tools.

### The Generalizability Gap

The observed performance decrements on independent validation align with broader patterns in medical imaging, where deep learning models frequently show degraded performance when applied to data from different sources than their training sets (Guan & Liu, 2022; Zech et al., 2018). In optic nerve histology, potential sources of domain shift include differences in tissue preparation protocols, staining intensity and consistency, image acquisition parameters, and anatomical variation across species and strains.

The magnitude of the generalizability gap varied by model, suggesting that certain architectural or training choices may confer greater robustness. AxoNet 2.0 exhibited the smallest performance decrement (0.07 correlation points), possibly reflecting its more extensive training data or refined augmentation strategies. AxoNet showed the largest decrement (0.18 points), despite having been trained on rat tissue similar to our validation dataset. Such a difference may be attributed to AxoNet’s density estimation model, which may systematically fail to recognize small axons even with high correlation. This finding demonstrates that even same-species validation cannot guarantee generalizability across laboratories.

### Segmentation Quality versus Count Agreement

A notable finding from independent validation was the dissociation between segmentation quality and count agreement. Despite achieving moderate correlations for axon counts (r = 0.79 to 0.89), segmentation metrics revealed substantial limitations. Dice coefficients of 0.29 to 0.40 fell well below the published benchmark of 0.81, driven primarily by low recall values (0.18 to 0.27). This pattern, characterized by high precision (>0.94) and low recall, indicates conservative segmentation: models correctly identified axon pixels when they did so but missed substantial portions of actual axon area. This conservative segmentation pattern suggests models may be under-detecting axon boundaries or missing smaller axon profiles. For applications focused solely on axon counting, this may be acceptable if under-segmentation is consistent across images. However, for morphometric analyses requiring accurate axon size measurements, this conservative bias could systematically underestimate axon diameters and areas.

### Performance in Context

Despite the observed generalizability gap, the independent validation correlations of 0.79 to 0.89 remain potentially useful for many research applications. These values compare favorably to inter-observer variability reported for manual counting methods, which can exceed 10 to 15 percent coefficient of variation (Teixeira et al., 2014). The key consideration is whether the reduced accuracy is acceptable for specific experimental questions and whether systematic biases might confound treatment group comparisons.

The published Dice coefficients of 0.81 reported by both AxonDeep and AxoNet 2.0 warrant reconsideration in light of our independent validation findings. These values were obtained on held-out test sets drawn from the same data sources as training sets, representing within-distribution testing. Our observed Dice coefficients of 0.29 to 0.40 likely better reflect expected performance in typical research settings where models encounter novel tissue preparations.

### Limitations

Several limitations should be considered when interpreting these findings. First, our independent validation used a single dataset from one laboratory, limiting generalizability of the validation results themselves. Performance on other independent datasets may differ. Second, the validation dataset was limited to rat optic nerve tissue with PPD staining; performance on other species or staining methods was not assessed. Third, we evaluated models using their published implementations without fine-tuning, representing out-of-the-box performance that could potentially be improved with dataset-specific adaptation. Fourth, because AxonDeep (Deng et al., 2021) is not publicly available, we could not directly validate this model and instead tested AxonDeepSeg (Zaimi et al., 2018) as a publicly available alternative, which is an independently developed tool from a different research group with a different architecture and training paradigm. AxonDeepSeg results should therefore be interpreted as an evaluation of that tool’s applicability to optic nerve histology, not as a proxy for AxonDeep performance.

The scoping review itself has limitations. All included studies originated from a small number of research groups, potentially limiting diversity of approaches. The absence of studies applying machine learning to human optic nerve histology limits clinical translation. Heterogeneity in outcome reporting prevented direct meta-analytic synthesis.

### Future Directions

Several developments would strengthen the evidence base for machine learning in optic nerve histology. Multi-center validation studies using standardized datasets are needed to assess generalizability across laboratories and establish realistic performance expectations. Shared benchmark datasets with expert-consensus annotations would enable direct comparison of models and support development of domain adaptation techniques.

Standardization of reporting practices would improve comparison across studies. We recommend that future studies report correlation coefficients, mean absolute error, and Dice coefficients using consistent definitions, along with sample sizes and confidence intervals. Validation on held-out data from external sources should become standard practice before model publication.

Investigation of domain adaptation and transfer learning approaches may help bridge the generalizability gap. Techniques such as few-shot learning, unsupervised domain adaptation, and self-supervised pretraining have shown promise in other medical imaging domains and warrant exploration for optic nerve histology. Public release of model implementations would also facilitate independent validation and broader adoption.

## CONCLUSIONS

Current deep learning models for optic nerve axon histology achieve strong agreement with expert reference counts in within-study evaluations, with published correlation coefficients exceeding 0.96. However, independent validation reveals meaningful performance decrements when models are applied to novel datasets, with correlations ranging from 0.79 to 0.89 and segmentation Dice coefficients of 0.29 to 0.40. This generalizability gap demonstrates the importance of external validation before widespread adoption. Among tested models, AxoNet demonstrated the most robust performance on independent validation. Future work should prioritize multi-center validation studies, standardized benchmark datasets, public release of model implementations, and development of domain adaptation techniques to improve model generalizability across laboratories and tissue preparations.

## ACKNOWLEDGMENTS

The authors thank Shelby Graham, Kyle Freeman, Sophie Pilkinton, Andrew B. Stiemke, and William Edwards for their contributions in annotation of optic nerve cross-section images used in the independent validation dataset.

## AUTHOR CONTRIBUTIONS

**BC:** Conceptualization, Methodology, Validation, Software, Formal analysis, Visualization, Writing - original draft, Writing - review & editing. **NE**: Investigation, Writing - review & editing. **MYK**: Visualization, Writing – review & editing. **ND**: Investigation, Writing - review & editing. **JH**: Investigation, Writing - review & editing. **ZZ**: Investigation, Writing - review & editing. **GW**: Investigation, Writing - review & editing. **AP**: Investigation, Writing - review & editing. **HC**: Investigation, Writing - review & editing. **TJH:** Methodology, Resources, Investigation, Writing - review & editing. **MMJ:** Conceptualization, Supervision, Project administration, Funding acquisition, Writing - review & editing.

## DATA AVAILABILITY

A detailed summary of study characteristics and reported performance metrics is available as Table S3. Analysis code, and independent validation dataset supporting this review are available from the corresponding author upon reasonable request. Model implementations used for independent validation are publicly available from their original publications (AxoNet, AxoNet 2.0, and AxonDeepSeg).

