## Supplementary Materials for "Comparison of Deep Learning Tools for Optic Nerve Axon Quantification Finds Limited Generalizability on Independent Validation"

**Supplementary Figure S1. Detailed Comparison of Published vs. Independent Validation Performance for Optic Nerve Axon Quantification Models.** Axon count agreement between model predictions and manual reference counts is shown as Pearson coefficients (r) across all tested species. Blue bars indicate published benchmark values from original model publications, and red bars indicate independent validation performance on rat optic nerve histology (n = 57 images). AxonDeepSeg is not shown because there is no reported correlation metrics for axon count agreement in optic nerve tissue with this model.

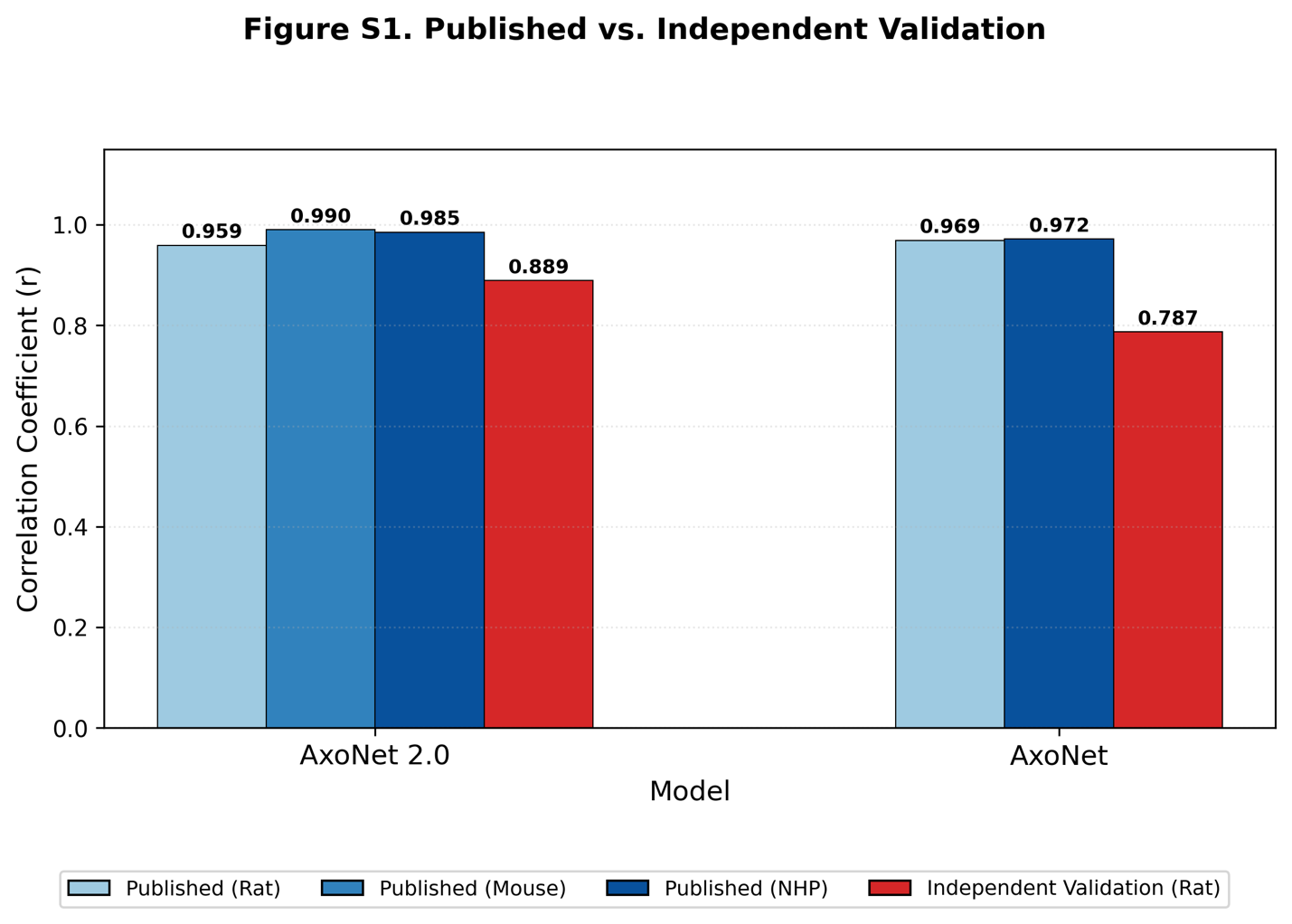

**Supplementary Table S1. Scoping Review Inclusion and Exclusion Criteria**

| **Category** | **Inclusion Criteria** | **Exclusion Criteria** |
| --- | --- | --- |
| Population / Sample | Studies involving human or animal optic nerve tissue, retinal ganglion cells, axons, myelin, astrocytes, microglia, oligodendrocytes, or fibroblasts | Studies excluding the optic nerve or related glial/axonal structures (e.g., purely cortical, spinal cord, or non-ocular tissues) |
| Intervention / Method | Use of machine learning (ML) methods for quantification, segmentation, morphometry, or histopathology analysis of optic nerve/glial tissue, where ML is understood to be any computational method that learns a mapping from input data to output predictions through optimization of model parameters | Studies limited to traditional histologic grading/manual counting without ML/DL components. |
| Outcomes | Reporting quantitative outcomes such as axon/myelin counts, morphometry, segmentation accuracy, detection performance, or comparative metrics (e.g., Dice score, sensitivity/specificity) | Studies without quantitative outcomes, or descriptive-only pathology papers without algorithmic analysis. |
| Study Type | Original research articles (basic science, translational, preclinical, or clinical) | Reviews, editorials, commentaries, conference abstracts without full data, non-peer-reviewed gray literature. |
| Language | English | Non-English |
| Publication Date | 2000-2025 (to capture modern ML/DL applications) | Studies published prior to 2000 (pre-modern ML) |

*Abbreviations*: DL = deep learning; ML = machine learning.

**Supplementary Table S2. Database Search Strategies**

| **Database** | **Search Strategy** |
| --- | --- |
| PubMed | ("optic nerve"[Title/Abstract] OR "retinal ganglion cell*"[Title/Abstract] OR axon*[Title/Abstract] OR myelin[Title/Abstract] OR astrocyte*[Title/Abstract] OR microglia*[Title/Abstract] OR oligodendrocyte*[Title/Abstract] OR glia[Title/Abstract] OR fibroblast*[Title/Abstract]) AND ("deep learning"[Title/Abstract] OR "machine learning"[Title/Abstract] OR "convolutional neural network"[Title/Abstract] OR CNN[Title/Abstract] OR segmentation[Title/Abstract] OR automated[Title/Abstract]) AND (histology[Title/Abstract] OR histopathology[Title/Abstract] OR histomorphometry[Title/Abstract] OR morphometry[Title/Abstract] OR quantification[Title/Abstract] OR grading[Title/Abstract]) |
| EMBASE | ('optic nerve':ti,ab OR 'retinal ganglion cell*':ti,ab OR axon*:ti,ab OR myelin:ti,ab OR astrocyte*:ti,ab OR microglia*:ti,ab OR oligodendrocyte*:ti,ab OR glia:ti,ab OR fibroblast*:ti,ab) AND ('deep learning':ti,ab OR 'machine learning':ti,ab OR 'convolutional neural network':ti,ab OR CNN:ti,ab OR segmentation:ti,ab OR automated:ti,ab) AND (histology:ti,ab OR histopathology:ti,ab OR histomorphometry:ti,ab OR morphometry:ti,ab OR quantification:ti,ab OR grading:ti,ab) |
| Scopus | (TITLE-ABS-KEY("optic nerve") OR TITLE-ABS-KEY(axon*) OR TITLE-ABS-KEY(myelin) OR TITLE-ABS-KEY(astrocyte*) OR TITLE-ABS-KEY(microglia*) OR TITLE-ABS-KEY(oligodendrocyte*) OR TITLE-ABS-KEY(glia) OR TITLE-ABS-KEY(fibroblast*) OR TITLE-ABS-KEY(retinal AND ganglion AND cell*)) AND (TITLE-ABS-KEY("deep learning") OR TITLE-ABS-KEY("machine learning") OR TITLE-ABS-KEY("convolutional neural network") OR TITLE-ABS-KEY(CNN) OR TITLE-ABS-KEY(segmentation) OR TITLE-ABS-KEY(automated)) AND (TITLE-ABS-KEY(histology) OR TITLE-ABS-KEY(histopathology) OR TITLE-ABS-KEY(histomorphometry) OR TITLE-ABS-KEY(morphometry) OR TITLE-ABS-KEY(quantification) OR TITLE-ABS-KEY(grading)) |
| Cochrane CENTRAL | ("optic nerve" OR "retinal ganglion cell*" OR axon* OR myelin OR astrocyte* OR microglia* OR oligodendrocyte* OR glia OR fibroblast*) AND ("deep learning" OR "machine learning" OR "convolutional neural network" OR CNN OR segmentation OR automated) AND (histology OR histopathology OR histomorphometry OR morphometry OR quantification OR grading) |

Searches conducted in January 2025. Date limits: 2000-2025. Language: English.

**Supplementary Table S3. Complete Data Extraction for Included Studies**

| **Study Characteristics** | | **Model Characteristics** | | **Dataset Characteristics** | | | | **Performance Metrics and Limitations** | | |
| --- | --- | --- | --- | --- | --- | --- | --- | --- | --- | --- |
| **Author/**  **Year** | **Institution** | **Model** | **Architecture/**  **Training** | **Optic Nerve Staining** | **Species** | **Strain/**  **Disease model** | **Sample Size** | **Reference Standard** | **Key Metrics** | **Limitations** |
| Ritch et al., 2020 | Georgia Institute of Technology; Emory University; Atlanta VA Healthcare System | AxoNet | U-Net encoder-decoder; outputs pixelwise axon count density estimates; Supervised | Toluidine blue | Rat | Brown Norway; Unilateral glaucoma induced by anterior-chamber injections | 27 ONs from 14 rats;  1,514 sub-images | Manual axon annotations by trained observers | R² = 0.938;  MAE = 4.4 axons/sub-image;  Bland–Altman LoA = ±14.3 axons | Trained on specific staining protocol; performance on other preparations unknown |
|  |  |  |  | PPD | NHP*  (rhesus and cynomolgus macaque) | Control and glaucoma eyes | ONs from 30 eyes;  494 sub-images |  | R² = 0.944;  MAE = 17.7 axons/sub-image;  Bland-Altman LoA =  [-43.9, 42.8] axons |  |
| Deng et al., 2021 | University of Iowa | AxonDeep | ResNeXt-50 encoder with feature pyramid decoder; semi-supervised GAN framework | PPD | Mouse | DBA/2J (various degrees of an inherited age-related form of glaucoma), D2.B6-*Lyst^bg-J^*/Andm (D2.*Lyst*; healthy), C57BL/6J (subjected to blast-induced TBI) and J:DO  (Diversity Outbred with healthy but natural variability from genetic background) | 56 ONs from 56 mice;  78 sub-images | Manual segmentations and center markings by expert annotators | R = 0.97;  Dice = 0.81 ± 0.04;  RMSE = 0.044 (normalized);  Mean absolute percent difference = 4.4% ± 3.3% | Validated only on mild-moderate damage; evaluated on subfields only; implementation not publicly available |
| Goyal et al., 2023a | Georgia Institute of Technology; Emory University; Atlanta VA; University of Iowa; Legacy Research Institute | AxoNet 2.0 | VGG16-based U-Net-style encoder-decoder CNN; soft Dice-loss optimization; Supervised | Toluidine blue | Rat | Brown Norway; unilateral ocular hypertension induced by anterior chamber injection | 46 eyes from 23 rats;  1,421 sub-images | Pixelwise annotations by two masked human annotators combined into probabilistic maps | Rat: R² = 0.92;  Dice = 0.81 ± 0.02;  RMSE = 6.18 axons  Bland-Altman LoA=  [-6.84, 13.57] axons | Cannot reliably segment degenerating axons; brightfield resolution constraints; inter-lab variability in defining normal axons |
|  |  |  |  | PPD | Mouse** | DBA/2J; inherited glaucoma and blast wave-induced traumatic optic neuropathy | ONs from 22 mice;  22 sub-images |  | R² = 0.98;  RMSE = 63.16;  Bland-Altman LoA =  [-115.4, 4.83] |  |
|  |  |  |  |  | NHPs*  (rhesus and cynomolgus macaque) | Glaucoma induced unilaterally by trabecular network laser treatment | ONs from 25 NHPs;  50 sub-images |  | R² = 0.97;  RMSE = 74.71;  Bland-Altman LoA = [7.95, 130.87] |  |
| Goyal et al., 2023b | Georgia Institute of Technology; Emory University; Atlanta VA Healthcare System |  | VGG16-based U-Net-style encoder-decoder CNN; soft Dice-loss optimization; Supervised; soft Dice loss optimization | Toluidine blue | Rat | Brown Norway; Unilateral ocular hypertension induced by anterior chamber injection | Cohort IV: 9 ONs from 9 normotensive eyes;  Cohort V: 68 eyes from 34 rats with unilateral ocular hypertension;  Cohort VI: 33 eyes from 23 rats with unilateral ocular hypertension | Morrison grading scale (MGS) for semi-quantitative damage scoring | N/A | Cannot segment degenerating/  severely damaged axons; image stitching artifacts; border inaccuracies |

Summary of study characteristics, model specifications, and reported performance metrics extracted from the four manuscripts meeting inclusion criteria. All studies focused on deep learning approaches for automated quantification of retinal ganglion cell axons in optic nerve cross-sections from experimental models. Goyal et al., 2023a and 2023b describe the same model (AxoNet 2.0) in separate publications focusing on different aspects: 2023a emphasizes cross-species validation and morphometric capabilities, while 2023b focuses on its application to experimental glaucoma models. Goyal et al. 2023b does not report traditional model performance metrics. *Species not explicitly named in Ritch et al. (2020) and in Goyal et al. (2023b); NHP dataset were annotated from Reynaud et al. (2012) from rhesus and cynomolgus macaques from Burgoyne Lab (Devers Eye Institute). **Mouse dataset was annotated from Deng et al. (2021). *Abbreviations*: GAN = generative adversarial network; CNN =convolutional neural network; MAE = mean absolute error; MGS = Morrison grading scale; NHP = non-human primate; ON = optic nerve; RMSE = root mean squared error; TBI =traumatic brain injury; LoA = limits of agreement; PPD = p-paraphenylenediamine.
